# A High-quality Genome Assembly of *Annona squamosa* (Custard Apple) Provides Functional Insights into an Emerging Fruit Crop

**DOI:** 10.1101/2024.12.26.630397

**Authors:** Manohar S. Bisht, Shruti Mahajan, Abhisek Chakraborty, Vineet K. Sharma

## Abstract

*Annona squamosa*, also known as custard apple, is an emerging fruit crop with medicinal significance. We constructed a high-quality genome of *A. squamosa* along with transcriptome data to gain insights into its phylogeny, evolution and demographic history. The genome has an N50 value of 93.2 Mb assembled into seven pseudochromosomes. The demographic history showed a continuous decline in the effective population size of *A. squamosa*. Phylogenetic analysis revealed that magnoliids were sister to eudicots, whereas ASTRAL gene trees showed discordance due to incomplete lineage sorting (ILS). Genome syntenic and Ks distribution analyses confirmed the absence of a recent whole genome duplication event in the *A. squamosa*. Gene families related to photosynthesis, oxidative phosphorylation and plant thermogenesis were found to be highly expanded in the genome. Comparative analysis with other magnoliids revealed the adaptative evolution in the genes of flavonoid biosynthesis pathway, amino sugar, nucleotide sugar and sucrose metabolism, conferring medicinal value and enhanced hexose sugar accumulation. In addition, we performed genome-wide identification of *SWEET* genes. Our high-quality genome and evolutionary insights of this emerging fruit crop, thus, serve as a valuable resource for advancing studies in functional genomics, evolutionary biology, and crop improvement.

## Introduction

Fruit crops have been under cultivation for centuries and are an integral part of human life. Many fruiting crops have gained much importance in their breeding and domestication, like Apple (*Malus domesticus*), Mango (*Mangifera indica*), Grape (*Vitis* sp.), Banana (*Musa* sp.) etc. However, many fruit crops with commercial potential are currently classified as “emerging” fruit crops (Hummer et al. 2012; Wang et al. 2023). One such family of underutilised fruiting crops is the Annonaceae family, the largest family of the order Magnoliales, majorly distributed in tropical regions of the world (Datiles and Acevedo-Rodríguez 2022). Genus *Annona*, *Asimina*, *Rollinia*, and *Uvaria* are the four edible fruit-bearing genera in Annonaceae, among which *Annona* is the most important source of edible fruits, which consist of some important species of this genus (*Annona squamosa, Annona reticulata, Annona muricata, Annona cherimola,* and *Annona mucosa*), in which, *Annona squamosa* (custard apple) is the most widely cultivated species (Padmanabhan and Paliyath 2016).

*Annona squamosa* (2n=14) is a small deciduous tree (about 3-6 meters in height). The leaves are alternate, ovate or elliptic-oblong. Flowers are pendulous, fragrant and yellowish green with three sepals and three petals. The fruit is a syncarp and irregularly heart-shaped (Datiles and Acevedo-Rodríguez 2022) (**Figure 1 A-C**). *A. squamosa* is natively from the New World tropics and most widely cultivated *Annona* spp. in the tropical regions of Africa, America, Asia, and the Pacific (Datiles and Acevedo-Rodríguez 2022).

**Figure 1.**
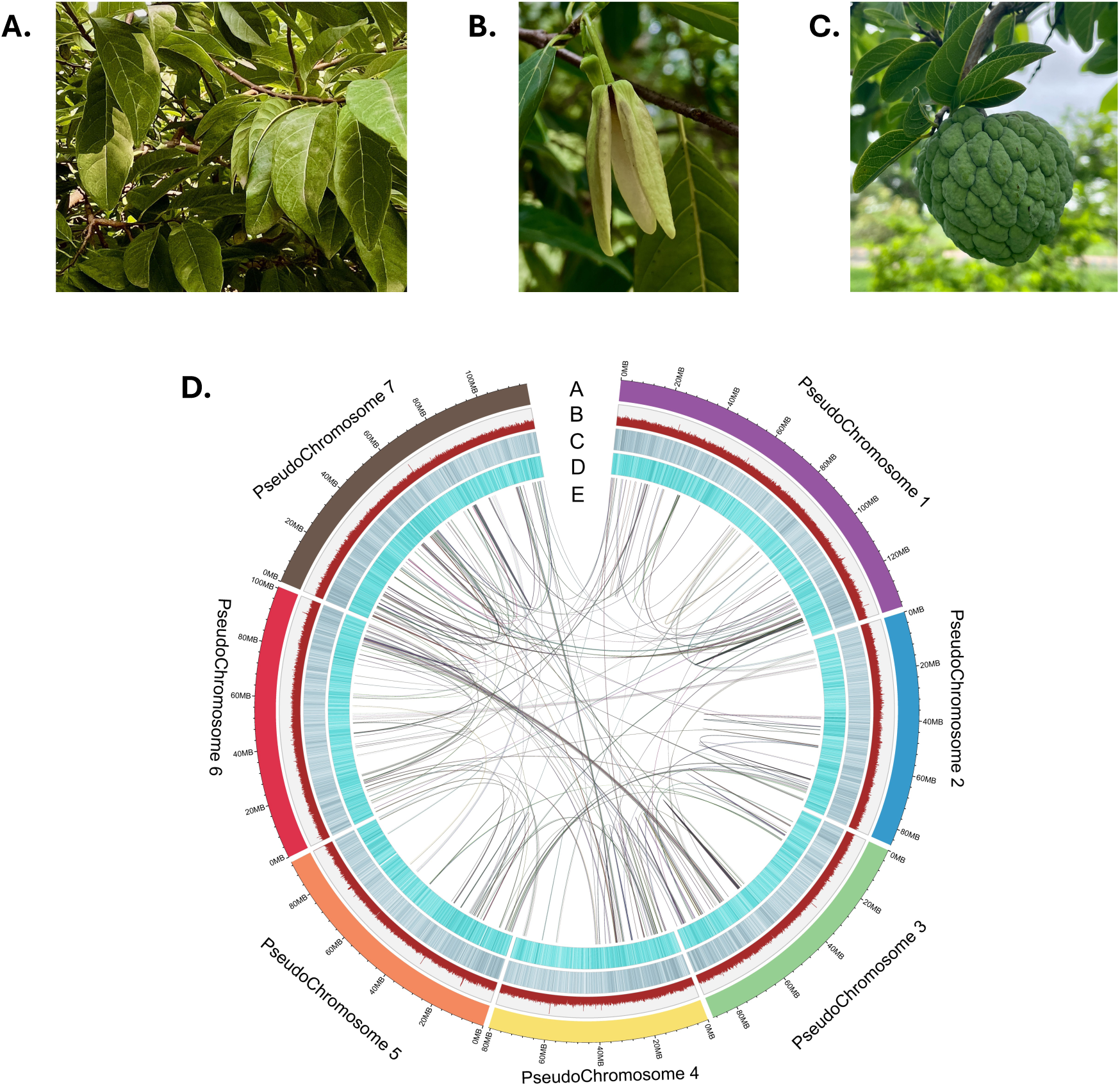
Morphology and genome assembly characteristics of *A. squamosa*. A. Leaves B. Flower C. Fruit D. Overview of genome assembly a. Assembled pseudochromosomes b. GC density c. Repeat density d. Gene density e. Chromosome synteny

The extracts from various parts of *A. squamosa*, such as roots, bark, leaves, and fruit, possess a wide range of ethnomedicinal importance conferring to its anticancer, antioxidant, antiparasitic, antimalarial, antidiabetic, insecticidal, microbicidal antihypertensive, hepatoprotective, and molluscicidal activities (Ma et al. 2017). Phytochemical analysis extracts from seed, leaf, bark, and roots reveal that the plant is filled with secondary metabolites like alkaloids, flavonoids, tannins, etc. (Kumar et al. 2021a). In addition to the immense medicinal properties, the fruit of *A. squamosa* is referred to as ‘one of the most delicious fruits known to man’ because of the high sugar content (∼28%) compared to other sweet fruits such as mango (∼15%) and banana (∼20%) (Forster et al.; Dar et al. 2016a). Further, among the other *Annona* species, *A. squamosa* has a higher total soluble sugar content (Anuragi et al. 2016). However, the genome of *A. squamosa*, the most widely cultivated species of the genus Annona, was not yet available (Gupta et al. 2015; Fang et al. 2020a). Therefore, in this study, to sequence the genome of *A. squamosa*, we used Oxford Nanopore Technology (ONT) and 10X technology and reported the leaf transcriptome. Further, we examined the demographic history of *A. squamosa*, exploring population dynamics over time, and investigated its phylogenetic placement providing context for its evolutionary relationships with related species. Furthermore, we analysed the evolutionary history of *A. squamosa* with a focus on gene family evolution, whole genome duplication (WGD) events and adaptive evolution by comparing it with other members of Magnoliales. This genome resource will support future studies on genetic diversity, adaptive traits, and the biosynthesis of bioactive compounds, contributing to a deeper understanding of the medicinal potential and evolutionary biology of *A. squamosa*.

## Materials and Methods

### Sample collection, species identification and DNA extraction

The leaves from *A. squamosa* were collected from IISER Bhopal campus, Bhopal, India (23.2599°N, 77.4126°E). The DNA extraction was performed using DNeasy Plant mini kit (Qiagen). The extracted DNA was used for amplifying *matK* DNA marker region for species identification (**Figure S1**). The amplicons were sequenced on Sanger sequencer, followed by aligning with NCBI nucleotide (nt) database using blastn. The species was identified as *A. squamosa* with 99.88% identity of *matK.* After species identification, high molecular weight DNA was extracted from the leaves sample. So, the extraction buffer previously used for DNA extraction from *A. muricata* was modified for *A. squamosa* DNA extraction (Lira-Ortiz et al. 2020). The extraction buffer [3% CTAB (Cetyl trimethyl ammonium bromide), 1M Tris-HCl, 0.7M NaCl, 0.05M EDTA (Ethylenediamine tetra acetic acid), 3% PVP-40, 1% β-mercaptoethanol (freshly added)] was pre-heated for 30 mins at 65°C. The leaves used were already frozen for a few days to reduce the amount of photosynthesis products. The leaves were homogenized in liquid nitrogen using a pre-cooled autoclaved mortar pestle. The powdered leaves were added to 1 mL of pre-heated extraction buffer and mixed by inversion. Proteinase K (25 µL) and RNase A (2 µL) were added for protein and RNA hydrolysis, respectively. The tubes were mixed by inversion and incubated for 12 hrs at 65°C. Upon lysis, the DNA was purified twice using chloroform: iso-amyl alcohol (24:1) in equal volume to remove the organic contaminants. The aqueous phase was taken and 0.7x ice-cold isopropanol was added to precipitate the DNA. An overnight incubation at -20 °C was done to facilitate DNA precipitation. After this incubation, the DNA was pelleted down and followed by washing with 70% ethanol. The washed DNA was dissolved in the G2 buffer of Blood and cell culture DNA mini kit (Qiagen) which was facilitated by incubation for 30 mins at 50°C. The G2 buffer with dissolved DNA was passed through equilibrated Genomic tip 20 and allowed to pass under gravitation pull. The Genomic tip 20 column was three times washed with 1 mL of QC buffer. The DNA was eluted in QF buffer (1mL). For DNA precipitation, 0.7x isopropanol was added and kept at -20°C for overnight. After incubation, the DNA was pelleted down, washed thrice with 70% ethanol, and air-dried to remove remnant ethanol. Finally, the pellet was suspended in 50 µL of nuclease-free water. The DNA sample was quantified on Qubit 2.0 fluorometer using a qubit ds DNA broad-range assay kit (Invitrogen, United States). The DNA samples were purified using 0.45x magnetic beads (Beckman Coulter, USA) for nanopore sequencing.

### Genome sequencing

The extracted DNA was utilised for linked read sequencing where it was used for preparation of a linked read library on the Chromium platform with the help of Chromium Genome Library Kit and Gel Bead Kit v2 (10x Genomics, USA). The linked read library was loaded on NovaSeq 6000 (Illumina, Inc., USA) for 150 bp paired-end sequencing. The library preparation for nanopore sequencing was done using ONT library preparation kits (SQK-LSK109 and SQK-LSK110). The libraries were sequenced on an in-house MinION Mk1C sequencer using FLO-MIN106 flowcells.

### RNA extraction and sequencing

The extraction buffer used for RNA extraction from *A. squamosa* leaves was the same as that used for *A. muricata* fruits (Montenegro Brasil et al. 2008). The protocol starts with the leaf homogenate, which was transferred into the RNA extraction buffer. The lysis was performed for 15 mins at 60°C. After incubation, the samples were cooled down on an ice bath for 10 mins. 150 µL of 5M Potassium acetate was added to cooled samples, mixed carefully, and centrifuged at 9,500 xg for 20 mins at room temperature. The supernatant was taken and mixed with 0.5 volume of 95% ethanol. The mixture was passed through the column of RNeasy mini kit (Qiagen). Further process was followed as per the manufacturer’s protocol. Qubit 2.0 fluorometer was used to quantify the RNA using RNA HS assay kit (Invitrogen, USA). The extracted RNA was used for library preparation using TruSeq stranded total RNA library preparation kit along with Ribo-Zero Plant workflow. The library quality assessment was performed on Agilent TapeStation 4150 with a High Sensitivity D1000 ScreenTape. The library was loaded on NovaSeq 6000 (Illumina, Inc., USA) and sequenced for generating paired-end reads of 150 bp.

### Genome size and ploidy estimation

The genome size of *A. squamosa* was computationally estimated by preprocessed 10x linked reads using Jellyfish v2.3.0 (Marçais and Kingsford 2011) that created K-mer count histograms, which were later employed in GenomeScope 2.0 (Ranallo-Benavidez et al. 2020) for genome size and heterozygosity estimation. Further, the ploidy estimation was performed using Smudgeplot v0.2.2 (Ranallo-Benavidez et al. 2020).

### Genome assembly and assessment

Oxford Nanopore sequencing raw reads were base-called using Guppy v3.2.1 (Oxford Nanopore Technologies), and adapters were removed using Porechop v0.2.4 (Oxford Nanopore Technologies). *De novo* assembly was performed with different genome assemblers using default parameters: Wtdbg2 (Ruan and Li 2019), Canu (Koren et al. 2017), and Flye v2.9.1 (Kolmogorov et al. 2019). Based on Quast v5.0.2 (Gurevich et al. 2013) statistics (N50, L50, assembled genome length and number of contigs), the Flye genome assembly was selected and polished using Pilon v1.23 (Walker et al. 2014) in three iterations. The scaffolding of contig assembly was performed by ARCS v1.2.2 (Yeo et al. 2018) and LINKS v2.0.0 (Warren et al. 2015) using barcode-filtered 10x linked reads, and adapter-free Nanopore reads, respectively. AGOUTI v0.3.3 (Zhang et al. 2016) was used to further scaffold the obtained assembly using trimmed and filtered paired-end transcriptome data produced from Trimmomatic v0.39 (Bolger et al. 2014) (assembly v01).

10x linked reads were assembled using Supernova v2.1.1 (Weisenfeld et al. 2017) with the “-maxreads” option set to 681 million paired-end reads and other default parameters. The obtained assembly was corrected using barcode-processed 10x reads by Tigmint v 1.2.6 (Jackman et al. 2018). Further, the assembly was scaffolded using ARCS v1.2.2 (Yeo et al. 2018) and LINKS v2.0.0 (Warren et al. 2015) with the input of Longranger basic linked reads and adapter-free Nanopore reads, respectively. Further, scaffolding was done by AGOUTI v0.3.3 (Zhang et al. 2016) using trimmed paired-end transcriptome reads (assembly v02).

Assembly v01 and assembly v02 were merged using Quickmerge v0.3 (Chakraborty et al. 2016) to produce a hybrid assembly. Gap-closing was done by LR_Gapcloser (Xu et al. 2018) in five iterations and TGS_Gapcloser v1.2.1 (Xu et al.) using Nanopore reads. In this gap-closed assembly, final polishing was done using Pilon v1.23 (Walker et al. 2014). The assembly was further length-based filtered with ≥ 5kbp cutoff (assembly v03). Further, to orient and align the scaffolds into the pseudochromosomes, chromosome-level assembly of *A. cherimola* was used using Chromosemble in Satsuma v2 (Grabherr et al. 2010; Chakraborty et al. 2023a) with default parameters. The obtained pseudochromosome-level assembly (assembly v04) was then gap-filled using TGS_Gapcloser v1.2.1 (Xu et al.) and polished with Pilon v1.23 (Walker et al. 2014).

To assess the quality of assembly quality, Quast v5.2.2 (Gurevich et al. 2013) and BUSCO v5.4.3 (Manni et al. 2021) with the embryohyta_odb10 dataset were used. To further assess the contiguity of the assembly, barcode-filtered 10x linked reads, nanopore raw reads, and quality filtered transcriptomic reads were mapped to the final draft assembly using MiniMap2 v2.17 (Li 2018), BWA-MEM v0.7.17 (Li and Durbin 2009), and HISAT2 v2.2.1 (Kim et al. 2019), respectively, and SAMtools v1.13 (Li et al. 2009) “flagstat” utility was used to calculate mapping percentage.

### Demographic history

To gain insights into the demographic history of *A. squamosa,* Pair-wise Sequentially Markovian Coalescent (PSMC) (Li and Durbin 2011) was used. The filtered 10x short reads were mapped to the genome using BWA-MEM (Li and Durbin 2009). SAMtools v1.13 (Li et al. 2009) “mpileup” and BCFtools v1.9 (Li and Barrett 2011) were used to form the consensus diploid sequence. Sites with quality scores <20 were filtered from consensus sequences. PSMC analysis was performed using the parameters “N30 -t5 -r5 -p4+25∗2 + 4+6” with 100 bootstrap values. A generation time of 15 years and a per-generation mutation rate of 7 × 10^−9^ were used to infer the demographic history (Collevatti et al. 2014; Strijk et al. 2021).

### Gene set construction and functional annotation

The pseudochromosome-level genome assembly *A. squamosa* was used to create the *de novo* repeat library using RepeatModeler v2.0.3 (Flynn et al. 2020). Further, these repeats were clustered to remove the redundant sequences using CD-HIT-EST v4.8.1 (Li and Godzik 2006), and the remaining unknown TEs were classified using TEclass2 (Bickmann et al. 2023) software with a probability threshold of >0.65 and resulting library was used to soft mask the repeats using RepeatMasker v4.1.2 (http://www.repeatmasker.org).

The soft-masked genome was used to construct the gene set using MAKER v3.01.04 (Cantarel et al. 2008) in three rounds. In the first round, the evidence-based gene was predicted, and a *de novo* transcriptome assembly was constructed using Trinity (Haas et al. 2013) with transcriptome data from our study and previous study (Liu et al. 2016). Further, protein sequences from other magnoliids species *(A. muricata*, *A. cherimola*, *L. chinense,* and *M. grandiflora*) were used. In the second and third rounds of MAKER *ab initio* gene prediction was performed using AUGUSTUS v3.3.3 (Stanke et al. 2006) and SNAP v1.0 (Korf 2004). The obtained gene models were then filtered for AED value <0.5 and transcript length ≥150 bp.

Additionally, Barrnap v0.9 (https://github.com/tseemann/barrnap), miRBase database (Kozomara et al. 2019) (sequence identity 80%, e-value 10-3) and tRNAscan v2.0.9 (Chan and Lowe 2019), for rRNA, miRNA, and tRNA predictions, respectively.

The protein coding gene set was functionally annotated by performing BLASTP against NCBI-nr, Pfam-A (Finn et al. 2014)and Swiss-Prot databases (Bairoch and Apweiler 2000) with an e-value of 10^-5^. Further, *A. squamosa* gene set, including the genes in highly expanded gene families and evolutionary signatures, were annotated using eggNOG-mapper v2.1.12 (Cantalapiedra et al. 2021) and KAAS v2.1 (Moriya et al. 2007)

### Phylogeny construction and analysis of evolution in gene families

To resolve the phylogenetic position of *A. squamosa*, protein sequences of 23 plant species along with *A. squamosa* proteins were selected, which include eight Magnoliids (*Magnolia grandiflora, Liriodendron chinense*, *Annona muricata*, *Annona montana*, *Annona cherimola*, *Laurelia sempervirens*, *Aristolochia fimbriata*, *Piper nigrum*), six Eudicots (*Cynara cardunculus*, *Nicotiana attenuata*, *Glycine max*, *Manihot esculenta, Arabidopsis thaliana*, *Vitis vinifera*), six Monocots (*Dioscorea rotundata*, *Brachypodium distachyon, Musa acuminata*, *Oryza sativa*, *Sorghum bicolor*, *Zea mays*), two ANA-grade angiosperms (*Amborella trichopoda, Nymphaea colorata*) and one Gymnosperm as outgroup (*Ginkgo biloba*) (**Table S3**). For each protein, gene family clustering was performed on its longest isoforms using OrthoFinder v2.5.4 (Emms and Kelly 2019). The obtained orthogroups were further filtered to extract the fuzzy one-to-one orthogroups using KinFin v1.1 (Laetsch and Blaxter 2017). The resulting orthogroups were aligned using MAFFT v7.310 (Katoh and Standley 2013). BeforePhylo v0.9.0 (https://github.com/qiyunzhu/BeforePhylo) was used to filter and concatenate the alignments. A maximum-likelihood tree was constructed using RAxML v8.2.12 (Stamatakis 2014) with 500 bootstrap values and “PROTGAMMAAUTO” substitution model. In addition, the coalescence-based species tree with 5,087 gene trees was constructed using ASTRAL-III v5.7.3 (Zhang et al. 2018).

To examine the evolution in gene families, proteome files containing the longest protein isoforms were analysed using CAFÉ v5 (Mendes et al. 2021). As suggested for CAFÉ analysis, all vs all BLASTP was performed to cluster and filtering of gene families. Using the 163 million-year calibration period between *A. squamosa* and *N. attenuata*, the phylogenetic tree was transformed into an ultrametric tree (https://timetree.org/). The CAFÉ v5 with two-lambda (λ) model was performed using the ultrametric tree and filtered gene, in which species from Annonaceae family were assigned separated λ-value compared to other species. In addition, the protein sequences of five *Annona* species were used to find core gene family clusters and species-specific gene clusters using OrthoVenn3 (Sun et al. 2023).

### Genome synteny and whole-genome duplication (WGD) analysis

Synteny analysis was performed for inter-species (*A. squamosa – A. cherimola*) and intra-species (*A. squamosa-A. squamosa*) using MCScanX (Wang et al. 2012). Further, to infer WGD history in *A. squamosa* genome, whole genome Ks (substitutions per synonymous site) distribution was performed for paralogs and orthologs including *A. cherimola, A. montana,* and *L. chinense* using wgd2 (Chen et al. 2024).

### Adaptive evolutionary signatures in *A. squamosa*

For comprehensive insights into the adaptive evolutionary traits in *A. squamosa* genome, comparative evolutionary analysis was performed with 10 other magnoliids species, in which six species were from Magnoliales order (*A. montana*, *A. muricata*, *A. cherimola, M. biondii, M. grandiflora, L. chinense*), two from Piperales (*P. nigrum*, *A. fimbriata*) two from Laurales (*L. sempervirens*, *C. kanehirae*).

#### A. squamosa genes with unique substitution with functional impact

The orthogroups were constructed from the protein sequence of these 11 species using OrthoFinder v2.5.4 (Emms and Kelly 2019). Each orthogroup was filtered to keep the longest sequence per species, which was later aligned using MAFFT v7.310 (Katoh and Standley 2013). The genes of *A. squamosa* that had different amino acid in position as compared to other species were called genes with unique amino acid substitution. Gaps and ten positions around the gap were excluded from the analysis. Further, to screen the functional impact of these uniquely substituted genes, SIFT was used with UniProt as a reference database (Ng and Henikoff 2003).

#### A. squamosa genes with positive selection

The orthogroups nucleotide sequences of all 11 species were aligned using MAFFT v7.310 (Katoh and Standley 2013). The resulting PHYLIP format alignments with the phylogenetic tree of 11 species were used for the positive selection analysis by implementing the branch-site model in the “codeml” program of PAML v4.10.6 (Yang 2007). For the statistical significance, a likelihood-ratio test and chi-square analysis were performed, and genes that qualified the threshold against the null model with FDR-corrected (p <0.05) were named positively selected genes. These positively selected genes were then further identified for the positively selected codon sites (>95% probability) for the foreground lineage using Bayes empirical Bayes (BEB) analysis (Mahajan et al. 2023; Bisht et al. 2024).

### Genome-wide Identification of *SWEET* genes and phylogenetic analysis

Protein sequences for *SWEET* (sugars will eventually be exported transporter) gene were identified in the proteome of *A. squamosa* and other two *Annona* species (*A. montana* and *A. cherimola*), using the hidden Markov model (HMM) profiles of the MtN3_slv domain for the *SWEET* gene family (PF03083) which were downloaded from the Pfam database (http://pfam.xfam.org/) and used as query in HMMER software with e-value 10^-5^. All the resulting putative genes were then screened for the presence of the MtN3_slv domain using SMART (http://smart.embl-heidelberg.de/), and sequences with at least one MtN3_slv domain were retained.

To investigate and categorise these SWEET genes, the TAIR database (https://www.arabidopsis.org/) was used to retrieve the *Arabidopsis thaliana* AtSWEET proteins. MAFFT v7.310 (Katoh and Standley 2013) was used to align the genes, and BeforePhylo v0.9.0 (https://github.com/qiyunzhu/BeforePhylo) was used to filter and concatenate the alignments. RAxML v8.2.12 with “PROTGAMMAAUTO” substitution model and 500 bootstrap values was used to create a maximum-likelihood tree (Stamatakis 2014).

## Results

### Sequencing summary, genome size, ploidy, and heterozygosity estimation

DNA sequencing generated 111.8 Gb (150x coverage) using 10x linked reads sequenced on Illumina platform and 16.5 Gb (22x coverage) data from Nanopore sequencing. Transcriptome sequencing generated 15.8 Gb of raw short-reads data from the leaf tissues. The estimated genome size was 737.7 Mbp, with diploid ploidy and 0.18% heterozygosity.

### Genome assembly construction and quality assessment

The final pseudochromosome-level genome assembly had a genome size of 737.4 Mbp with N50 of 93.24 Mbp and GC of 34.32% (**Figure 1D**, **Table 1**). The assembly showed 94.1% complete BUSCO indicating its completeness. Further, 91.05% of the barcode-filtered linked reads, 91.33% of nanopore reads, and 95.57% of transcriptome data mapped to the final genome assembly.

**Table 1:** Genome assembly and annotation statistics of *Annona squamosa*.

| Feature | *A. squamosa* |
| --- | --- |
| **Final pseudochromosome assembly** |  |
| Total length (bp) | 730,367,479 |
| Number of scaffolds | 58 |
| Longest scaffold (Mb) | 137.68 |
| Scaffold N50 (Mb) | 93.21 |
| Scaffold L50 | 4 |
| Scaffold GC content | 34.32 % |
| BUSCO (complete) | 94.1 % |
| **Gene models** |  |
| Number of coding genes obtained from MAKER | 48,633 |
| Number of coding genes with AED value < 0.5 length-based filtering (High confidence gene set) | 40,435 |
| Repeat content of the genome (%) | 64.82 |
| **Non-protein-coding genes** |  |
| Number of rRNAs | 491 |
| Number of tRNAs (decoding standard amino acids) | 1,076 |
| Number of miRNAs (homology-based) | 95 |

### Gene set construction and functional annotation

A total of 64.82% of repetitive regions were identified in the *A. squamosa* genome, with 52.13% as interspersed repeats. Retroelements consisted of 29.96% LTR (long terminal repeat) elements (12.98% Ty1/Copia and 19.00% Gypsy/DIRS1 elements). MAKER predicted 48,633 protein-coding genes. After AED value-based (<0.5) and length-base (≥150 bp) filtering, 40,435 high-confidence genes were retained, and overall, 88% of these genes were functionally annotated against the publicly available databases. Further, these genes showed 88.3% of complete and fragmented BUSCO. Moreover, non-coding RNAs were also identified in *A. squamosa* genome assembly (**Table 1**).

### Demographic History of *A. squamosa*

PSMC results indicated a continuous decline in the effective population size of *A. squamosa*, correlating with its low heterozygosity of ∼ 0.18% (**Figure 2**). This might link to the Quaternary period’s contraction of tropical regions, suggesting that *A. squamosa* faced significant challenges from climate changes, like many other tropical species (Barlow et al. 2018; Strijk et al. 2021). In addition, *A. squamosa* might have experienced pronounced genetic bottlenecks during its domestication, further contributing to its reduced genetic diversity (Doebley et al. 2006; Zhu et al. 2007; Strijk et al. 2021).

**Figure 2.**
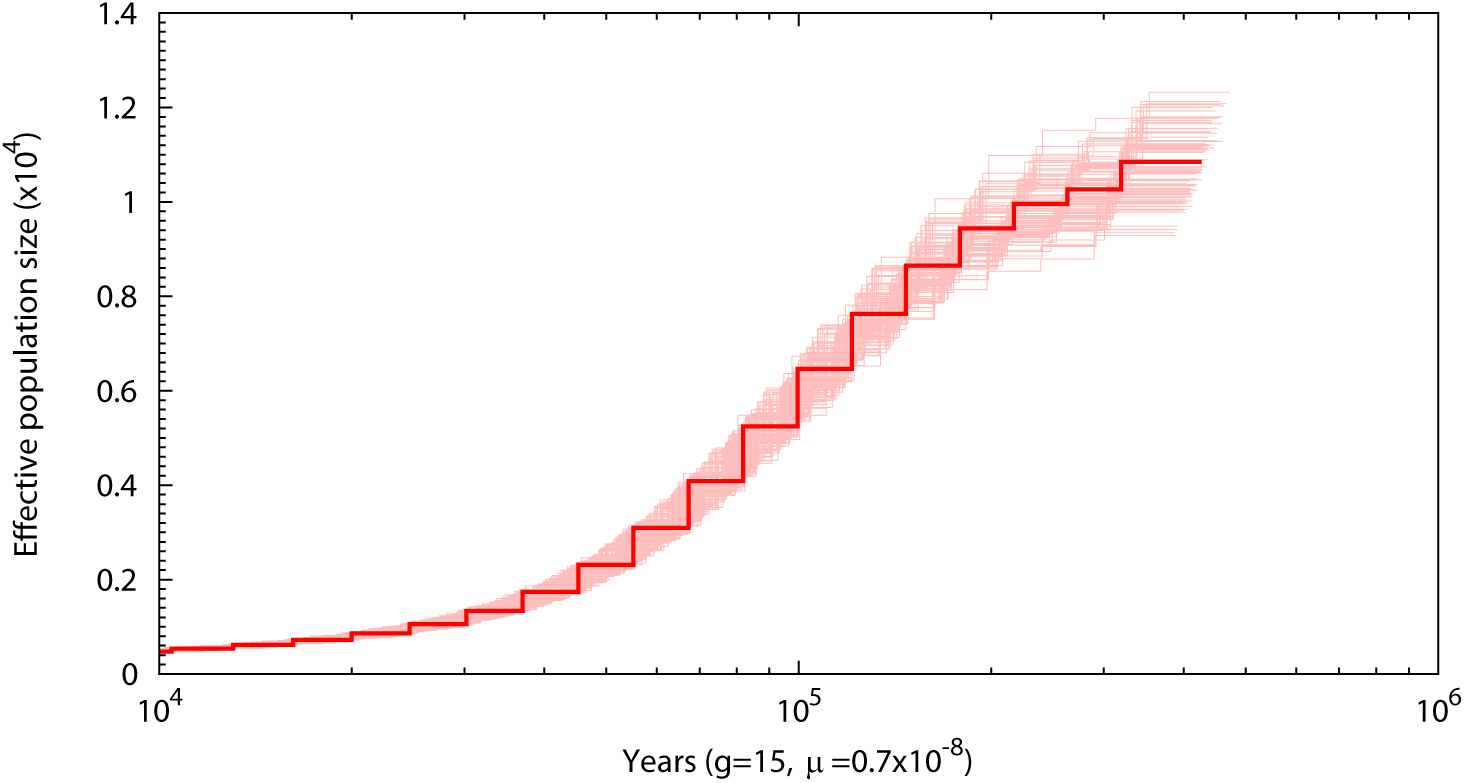
Demographic history of *A. squamosa*. Light red lines represent the bootstrap values used in PSMC analysis

### Phylogeny construction and gene family evolution

The Maximum Likelihood (ML) tree based on 433 one-to-one fuzzy orthogroups with 500 bootstrap replicates across 23 angiosperms and one gymnosperm (outgroup) revealed that *A. squamosa* was most closely related to *A. cherimola,* which was sister to the subclade formed by *A. montana* (Annonaceae). Additionally, magnoliids were positioned as sisters to eudicots (**Figure 3A**). In contrast, an ASTRAL concatenated amino acid tree based on 5,087 genes produced a different topology, placing magnoliids as a sister group to the eudicot-monocot clade. This discordance between the species tree and gene tree reflects the incomplete lineage sorting due to the rapid divergence of common ancestors of magnoliids, eudicots and monocots, as observed in other studies (Chen et al. 2018; Lv et al. 2020; Tang et al. 2023; Zhou et al. 2023).

**Figure 3.**
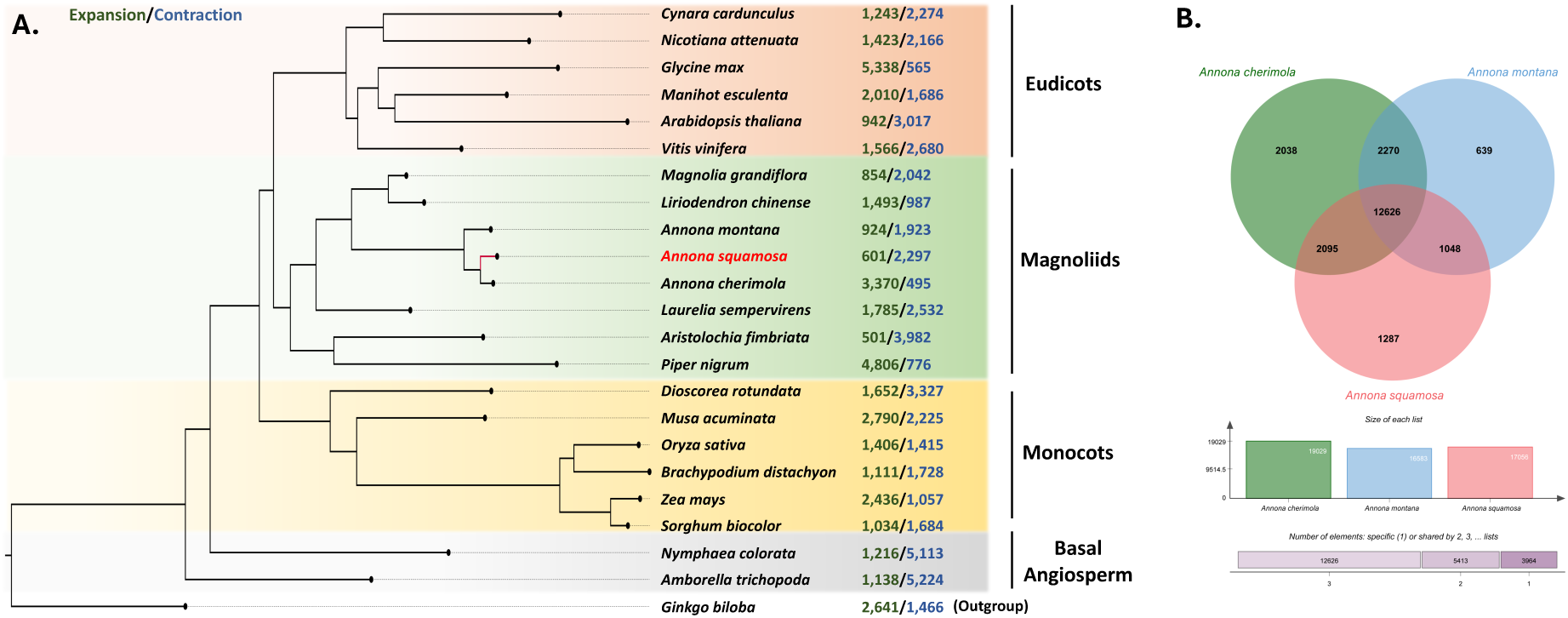
**. Genome-wide species phylogenetic tree of *A. squamosa* with respect to eudicots and monocots.** Numbers in green and blue represent the number of expanded and contracted gene families in each species, respectively. **B. Orthologous gene family cluster analysis of among *Annona* species**

Gene family evolution analysis identified a total of 9,443 filtered gene families, among which 1,204 gene families were expanded, and 1,506 gene families were contracted in *A. squamosa*. Further, among these expanded gene families in *A. squamosa*, 17 were highly expanded (>10 expanded genes). These highly expanded gene families were involved in photosynthesis, oxidative phosphorylation, biosynthesis of secondary metabolites, and thermogenesis etc.

Gene clustering among *A. squamosa* and four other *Annona* species showed 9,806 core gene family clusters and 1,035 species-specific gene clusters in *A. squamosa* (**Figure 3B)**.

### Whole-genome duplication and synteny

The Ks distribution of orthologs revealed *A. squamosa* to be diverged from *L. chinense* at around Ks ∼ 0.75, whereas *A. squamosa* diverged from *A. montana* at Ks ∼ 0.14 and from *A. cherimola* around Ks ∼ 0.04 (**Figure 4A**), which is congruent with species phylogeny (**Figure 3A**). Further, the paranome Ks distribution revealed no signs of independent WGD in *Annona* species after diverging. However, *L. chinense* experienced an independent WGD event at Ks ∼0.69 after diverging from *Annona* (**Figure 4B**) (Chen et al. 2018; Tang et al. 2023).

**Figure 4.**
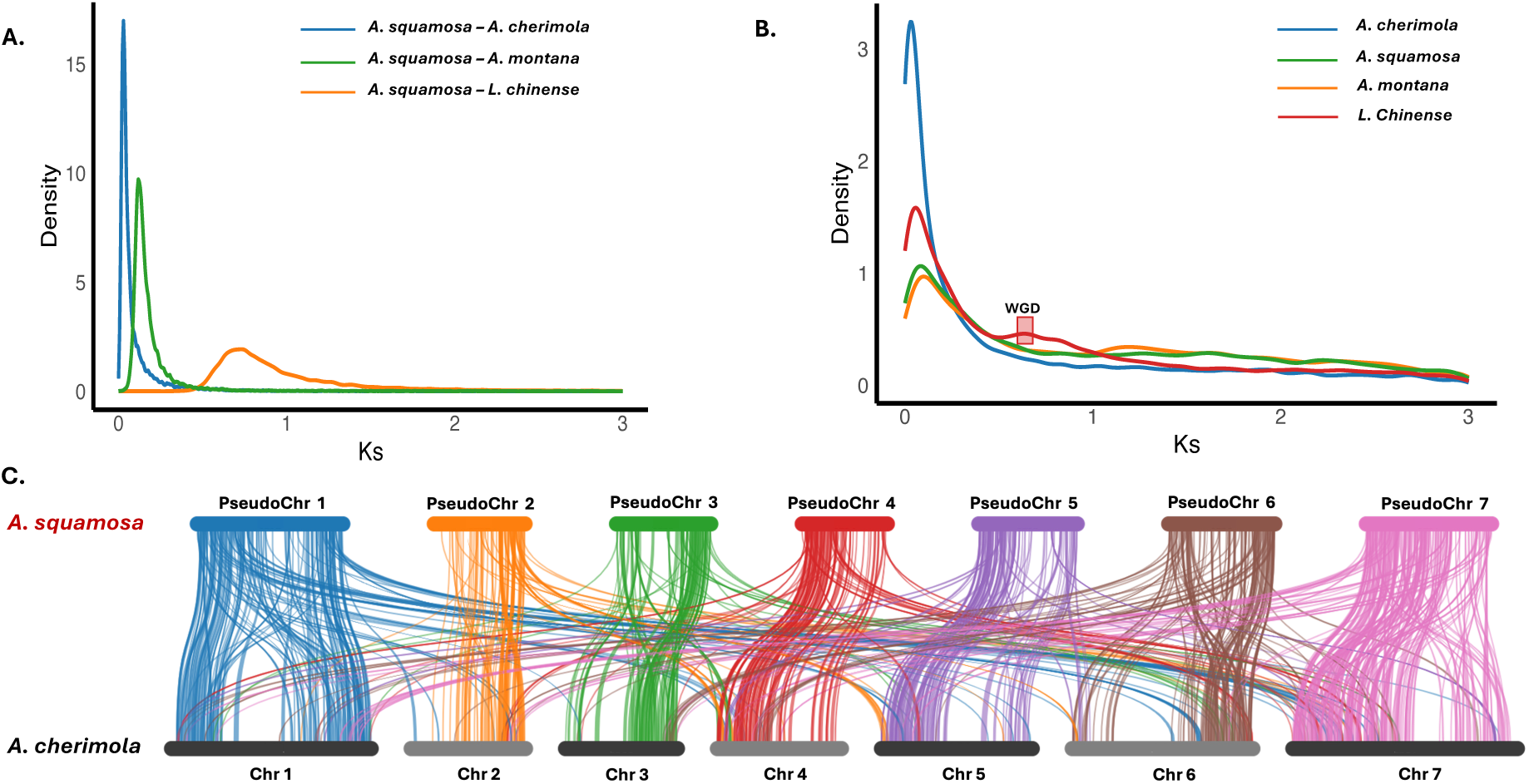
Whole genome duplication history of *A. squamosa*. A. Ks distribution of paralogs B. Ks distribution of orthologs C. whole-genome synteny between *A. squamosa* and *A. cherimola*.

The intra-genomic synteny of *A. squamosa* showed 9.30% of collinearity, while the inter-genomic synteny between *A. squamosa* and *A. cherimola* showed 35.21% of collinearity. This high level of interspecies synteny indicates a significant similarity between the genomes of the two species, reflecting their relatively recent divergence. In general, the majority of the chromosomes of *A. squamosa* were aligned with corresponding regions of the chromosomes of *A. cherimola* in a one-to-one relationship (**Figure 4C)**.

### Genes with adaptive evolution signatures in *A. squamosa*

In total, 1,781 orthogroups were found across 11 selected species of magnoliid clade to screen the genes with signatures of adaptive evolution in *A. squamosa*. Comparative analysis revealed 260 genes with unique amino acid substitution with functional impact and 75 positively selected genes (*p*-value <0.05) in *A. squamosa*. *Flavonoid biosynthesis pathway* Flavonoids are structurally diverse secondary metabolites in plants that contribute significantly to plant growth and development (Chakraborty et al. 2023b; Mahajan et al. 2024). Further, various phytochemical studies on *A. squamosa* revealed that various flavonoids, including rutin, kaempferol, quercetin, isorhamnetin, and farmarixetin, were rich in the leaves of plants in *Annona*, governing the medicinal importance (Gupta et al. 2008; Kumar et al. 2021b). In *A. squamosa,* genes involved in the biosynthesis of various flavonoids, including anthocyanins, showed evolutionary signatures. Further, these genes were found to have a high TPM value (>1) in the leaf tissues.

#### Amino sugar, nucleotide sugar and sucrose metabolism

*A. squamosa* fruit is one of the sweetest fruits, containing up to 28% of total sugars(Dar et al. 2016b; Fang et al. 2020b). Comparative evolutionary analysis revealed the adaptive evolution in the genes involved in amino sugar, nucleotide sugar and sucrose metabolism pathways. These evolutionary modifications in the genome of *A. squamosa* likely contribute to the metabolic efficiency in facilitating high sugar content and enhancing the plant’s adaptive responses to environmental pressures.

### Genes encoding *SWEET* transporters

The SWEET family is a novel class of sugar transporters that enable the diffusion of sugars along a concentration gradient across cell membranes by acting as bidirectional uniporters (Breia et al. 2021; Ji et al. 2022). A total of 13, 20, and 14 *SWEET* genes were Identified in the *A*. *squamosa, A. cherimola* and *A. montana* genomes, respectively. Phylogeny grouped these genes into four clades (**Figure 5**). Among 13 genes of *A. squamosa* eight genes were categorised into clades I and II, primarily transporting glucose. Three genes were in clade III, which mainly transports sucrose, and two genes were in clade IV, which transports fructose (Chen et al. 2012). Of these, clade I and IV genes were highly expressed in the leaf tissue compared to clade II and III, suggesting a key role of fructose and glucose accumulation over sucrose in the sink tissues.

**Figure 5.**
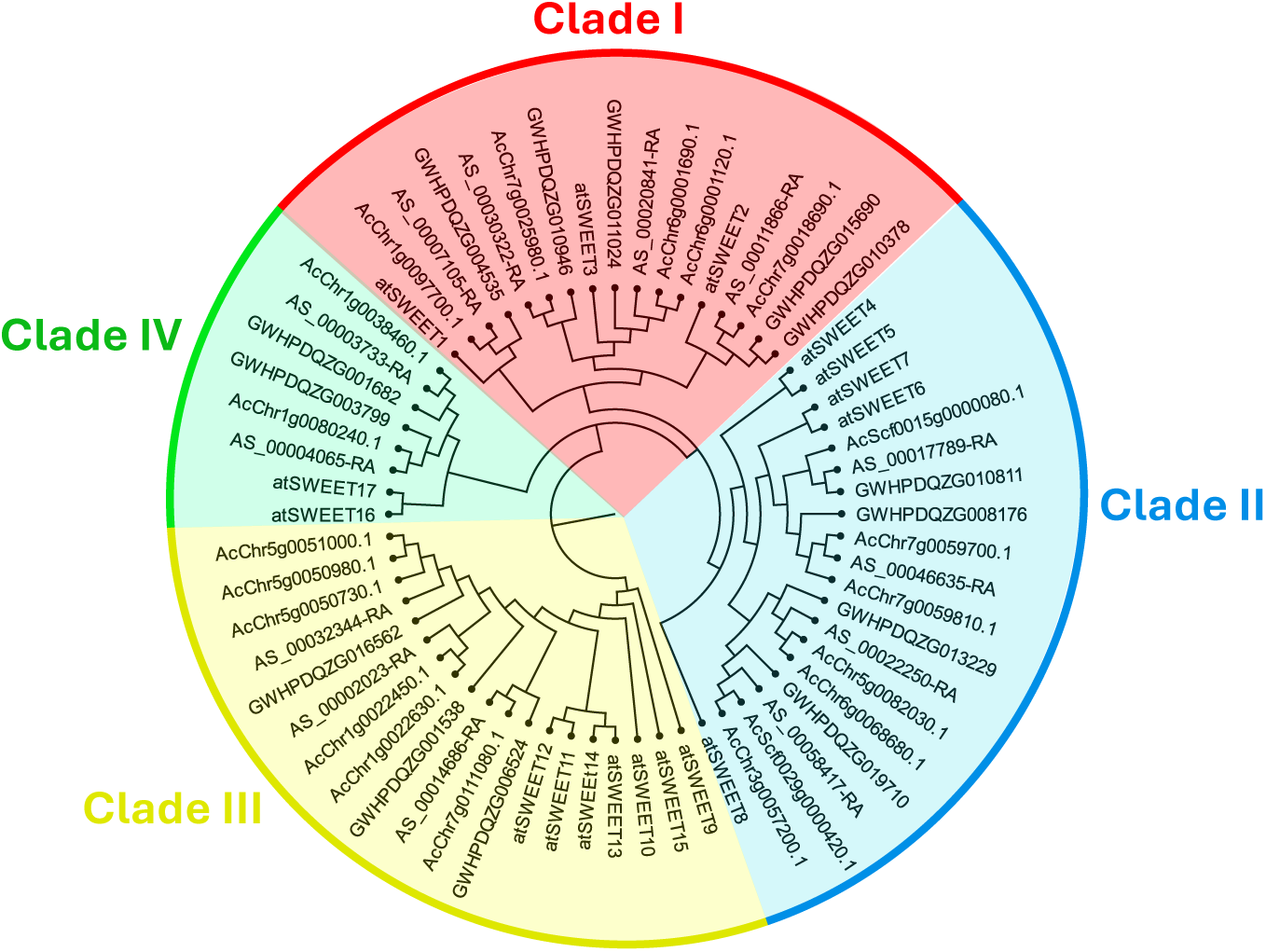
Phylogenetic relationship analysis of 64 SWEET proteins from *Annona squamosa*, *Annona cherimola*, *Annona montana*, and *Arabidopsis thaliana*.

## Discussion

Genome sequencing has advanced plant science, facilitating the production of numerous cash crops, including various fruit species. However, in the face of food security and sustainable agriculture, many fruiting crops are neglected and are being underutilised. The absence of genomic studies related to the genetic and functional diversity of these emerging crops creates a bottleneck in harnessing their full potential for improving yield, stress tolerance, and nutritional value.

In this study, we performed whole-genome sequencing of *A. squamosa*, which is an emerging fruit crop and provides the first high-quality genome of this species. The genome was diploid with an assembly size of 737 Mbp (**Table 1**). The heterozygosity of *A. squamosa* (0.18%) was low compared to 1.05% of *A. cherimola*, but higher than *A. muricata* (0.06%). This reduced heterozygosity of *A. squamosa* may result from extensive domestication and hybridization, as it is also one of the most widely cultivated species within the *Annona* genus, thus recognizing it as an emerging commercial fruit crop (Doebley et al. 2006; Zhu et al. 2007). Further, the low heterozygosity could also be a possible reason for the reduced effective population size (**Figure 2**), particularly in comparison to the higher effective population size observed in the more heterozygous *A. cherimola*. This decline in population size may raise challenges in the future in terms of the breeding of the plant (Zamir 2001; Doebley et al. 2006; Wang et al. 2023).

The rapid divergence of common ancestors of magnoliids, eudicots and monocots resulted in ancestral genetic polymorphisms, leading to incomplete lineage sorting (ILS) and discordance between species and gene trees (Rendón-Anaya et al. 2019; Qin et al. 2021). Our results, from the Maximum Likelihood (ML) tree based on 433 one-to-one fuzzy orthogroups and the ASTRAL tree derived from 5,087 genes, reflect this discordance (**Figure 3A)**. This incongruence is consistent with observations in other studies, which also report challenges in resolving ancient rapid radiations among early-diverging angiosperms due to ILS (Chen et al. 2018; Qin et al. 2021; Tang et al. 2023).

The pseudochromosome assembly of the *A. squamosa* genome provides a deeper understanding of the timing of the WGD event of the Annonaceae family. Our genome syntenic and Ks distribution analyses confirmed the absence of a recent WGD event in the *A. squamosa*, also observed in other *Annona* genomic studies (Tang et al. 2023; Talavera et al. 2023) (**Figures 4A and 4B**). This absence of WGD may suggest that the genus retained a more ancestral genome structure (**Figure 4C**).

Gene family evolution analysis reflects the evolutionary changes with respect to the ancestral species that are derived from various adaptive forces. These forces result in the expansion and contraction of gene families, resulting in lineage-specific adaptations that confer their distinct ecological and functional requirements (Luna and Chain 2021; Fang et al. 2022). The observed expansion of gene families involved in oxidative phosphorylation and thermogenesis in *A. squamosa* provides valuable insights into its floral biology. Thermogenesis facilitates the release of floral scents, serving as signals to attract insects by indicating the availability of food resources. (Kishore et al. 2012; Wang et al. 2014).

*A. squamosa* has been used in traditional medicines, governed by extracts obtained from various sections of the *A. squamosa* that contain phenol-based compounds, mainly alkaloids or flavonoids (Kumar et al. 2021b). The presence of evolutionary signatures in the enzymes involved in flavonoid biosynthesis suggested the adaptive evolution of these pathways in *A. squamosa.* Further, the high expression of these genes in *A. squamosa* leaf tissue indicates that these genes are actively transcribed, reflecting their functional importance, as leaves are the primary sites of flavonoid biosynthesis (Marinova et al. 2007; Falcone Ferreyra et al. 2012).

*A. squamosa* fruits are largely rich in reducing sugars (Glucose, fructose, etc.) compared to non-reducing sugar (sucrose etc.)(Brandão and Santos 2016; Fang et al. 2020b), contributing to their sweetness and shorter shelf life, as sucrose generally enhances fruit shelf life (Yu et al. 2022; Cardoso et al. 2023). Our genomic analysis revealed an adaptive evolution in genes associated with amino sugar, nucleotide sugar, and sucrose metabolism, suggesting a selective shift towards this high hexose sugar accumulation. In addition to that, the high expression of *SWEET* transporters genes related to clade I and IV over clade III suggest a preponderant role of fructose and glucose transport over sucrose transport in the sink tissues. This, high hexose-to-sucrose ratio in early fruit development has been reported to increase mitotic activity through hexokinase (*HK*) signalling that increases cell number. Additionally, hexoses have higher osmotic potential than sucrose, causing them to retain more water, which increases fruit size and cell volume—a property that affects shelf life, nutritional qualities, and consumer acceptance. (Yu et al. 2022).

## Conclusion

In summary, the high-quality genome of this emerging sweet fruit crop, along with evolutionary insights and pattern of gene family expansion, with phylogenetic relation, provides a deeper understanding of the adaptations that shape the evolution of *A. squamosa* within the Annonaceae family. This study, thus, serves as a valuable resource that will assist in the field of functional genomics, evolutionary biology, and crop improvement in the Annonaceae family.

## Acknowledgments

MSB thanks the Ministry of Education, Govt. of India, for the Prime Minister Research Fellowship (PMRF). S.M. and A.C. thank the Council of Scientific and Industrial Research (CSIR) for the fellowship. The authors also thank the Sanger sequencing at IISER Bhopal and the intramural research funds provided by IISER Bhopal.

## Author’s contribution

V.K.S. conceived and coordinated the project. S.M. performed DNA-RNA extraction, prepared the samples for sequencing, performed Nanopore sequencing, and species identification. M.S.B. and V.K.S. designed the computational framework of the study. M.S.B. performed all the computational analyses presented in the study with inputs from A.C. M.S.B. constructed all the figures. M.S.B. and V.K.S. interpreted the results. M.S.B., A.C., S.M., and V.K.S. wrote the manuscript. All the authors have read and approved the final version of the manuscript.

